# Pernicious liaisons: antibiotic-depressed immune response of a livestock micropredator, the common vampire bat (*Desmodus rotundus*)

**DOI:** 10.1101/2023.09.15.557909

**Authors:** Iván Cabrera-Campos, Rafael Ávila Flores, David Alfonso Rivera-Ruiz, L. Gerardo Herrera M.

## Abstract

Antibiotics are pharmaceutical products that have the potential to affect the immune performance of wildlife. Wildlife species might incorporate antibiotic residues in their system by feeding on livestock treated with these chemicals. One of the most important interactions of livestock with wildlife is that established with the common vampire bat (Desmodus rotundus). We used vampire bats as an ecologically relevant model to test the effect of antibiotics on wildlife immune response. We tested the effect of clindamycin on the humoral and cellular acquired immune responses of common vampire bats captured in the wild in southern Mexico. We expected that both cellular and humoral acquired immune responses would be negatively affected after bats were exposed to clindamycin for several days. We measured local inflammation and serum immunoglobulin concentration (IgG) after the repeated application of phytohemagglutinin. We expected that antibiotic-exposed bats would present a weaker inflammatory response to a second injection of PHA and that their IgG serum levels did not increase to the same rate after the third PHA injection.

Antibiotic-treated vampire bats exhibited weaker inflammatory response to the repeated PHA treatment: induced swelling was ∼30% larger after the second injection than that after the first injection, whereas swelling after the second injection in antibiotic-treated bats was ∼10% lower than after the first injection. There was an increase of IgG serum levels following three consecutive PHA injections but it occurred only in vampires that did not receive the antibiotic: IgG serum levels of control individuals increased ∼90% over pre-injection values, whereas this value was ∼15% lower in vampires treated with antibiotic. Our study adds to previous evidence pointing to the negative effect that exposure to anthropogenic chemicals generates in wildlife capacity to maintain a healthy immune system and warrants further work on the relationship of potential antibiotic-induced changes in gut microbiota and immune response.

## Introduction

The global presence of pharmaceuticals in the environment constitutes a major threat to wildlife [1, 2] with side effects ranging from organ failure [3], proliferation of antibiotic-resistant bacteria [4], and behavioral changes [5]. Antibiotics, defined as pharmaceutical products that destroy or slow down the growth of bacteria and their residues in the environment, might affect wildlife populations at several trophic levels [6] Exposure to antibiotic residues has the potential to affect the immune performance of wildlife as it has been shown in laboratory and farm animals. For instance, antibiotic treatment decreased immunoglobulin production in house mice (*Mus musculus* [7, 8]), common carps (*Cyprinus carpio* [9]) and domestic pigs (*Sus scrofa domestica* [10]), and hampered the cellular immune response of common carps [11].

One of the routes by which wildlife species might incorporate pharmaceutical residues in their system is by feeding on livestock, poultry or fish treated with these chemicals. For instance, the decline of vulture populations *Gyps bengalensis* in Pakistan was attributed to feeding on cattle treated with anti-inflammatory drugs [3]. In spite of recent policies to discourage their use, antibiotics are still widely used in the livestock industry in America [12]. One of the most important interactions of livestock with wildlife throughout America is that established with the common vampire bat *(Desmodus rotundus;* [13, 14]). The common vampire bat feeds on blood of several mammal species and it acts as a micropredator of livestock [15]. The potential impact of antibiotic exposure via blood ingestion has not been examined in vampire bats, but there is limited evidence suggesting relevant changes. For instance, antibiotic resistant bacteria were recently reported in common vampire bats suggesting possible cross-species exchange of bacteria with livestock [16, 17]. Therefore, the common vampire bat is an ecologically relevant model to test the effect of antibiotic exposure on the immune response of wildlife.

We tested the effect of clindamycin, a commonly used lincosamide antibiotic used to prevent bacterial replication by interfering with the synthesis of proteins, on the humoral and cellular acquired immune responses of common vampire bats captured in the wild in southern Mexico. We stimulated acquired cellular response after two applications of phytohemagglutinin (PHA), a proinflammatory protein extracted from the red bean (*Phaseollus vulgaris*). When injected in the skin, PHA stimulates the mitogenic activity of several immune cells and produces local inflammation [18]. The inflammation produced by a single application of PHA measures the strength of the individual’s innate immune response [19, 20]. However, the repeated application of PHA stimulates a stronger local inflammatory response increasing circulating lymphocyte levels and it is considered as an accurate measure of acquired T-cell-mediated immune response [21-23]. We evaluated changes in humoral acquired immune response measuring IgG serum levels after 3 repeated applications of PHA. We expected that both cellular and humoral acquired immune responses would be negatively affected after bats were exposed to clindamycin for several days. Accordingly, we expected that antibiotic-exposed bats would present a weaker inflammatory response to a second injection of PHA and that their IgG serum levels did not increase to the same rate after the third PHA injection.

## Methods

### Capture and management of individuals

We collected 29 non-reproductive adult males of *Desmodus rotundus* in February-July 2021 in three caves (17.526772, -92.732864; 17.536069, -92.757609; 17.549367, -92.756418) located in Poaná, Tabasco. The climate of the area is hot-humid, and the vegetation is fragmented tropical forest with large patches of grassland used for livestock [24]. The bats were captured from 19:30 to 06:00 hours with mist nets at the entrance or the cave or with butterfly nets inside the caves. We recorded forearm and total body length (± 0.01 mm), body mass (± 0.1 g), and sex of each individual and ringed with numbered bracelets for identification. Bats were transferred to a portable cage after capture to be transported to the experimental setting in the city of Villahermosa, Tabasco.

### Experimental procedure

Bats were individually kept in small cages (20 x 20 x 20 cm) placed on the top of a table inside a dark room located in the Parque Museo La Venta, in Villahermosa. Bats were randomly assigned to an experimental or a control group the day of their capture (day 0). The experimental group (n = 15) was fed 0.5 mg of clindamycin per ml of cow blood [25], and the control group (n = 14) was fed cow blood without the antibiotic. Room temperature (24-25°C) and humidity (70-80%) were kept relatively constant using and air conditioning system. Blood was served at the same time (10:00-11:00 local time) every day over the course of the experiment. Once the experimental trials were finished, individual bats were maintained permanently in captivity for conducting behavioral experiments in a parallel research project. Both field and captivity procedures followed the guidelines published by the American Society of Mammalogists [26] and were conducted under the research and scientific collecting license of the Mexican environmental authorities (permit number SGPA/DGVS/00792/21).

### Evaluation of the adaptive cellular response

Bats were injected with PHA on days 1, 10 and 15 after their capture. On each occasion, we injected 50 µL [27] of a 3 mg/mL solution of PHA (PHA-P, no. L8754, Sigma-Aldrich, Mexico) in phosphate-buffered saline (PBS, Sigma-Aldrich, Mexico [28]) in the right ankle of each individual. As a control, 50 µL of PBS (0.01 M) were injected in the left ankle. Average thickness (from three measures) of both ankles was measured with a digital caliper (Trupper, Mexico, ±0.01 mm) 10 min before and 6 and 12 h after PHA or PBS injection. PHA response was evaluated as the swelling index (SI) 6 and 12 h after the injection as

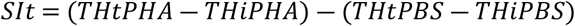

where *THt* is the ankle thickness 6 or 12 h after the injection and *THi* is the ankle thickness before the injection of PHA or PBS [29]. A *SI* value of 0 occurs when swelling in the PHA-injected foot is equal to that in the PBS-injected foot and indicates no or low immune response. We found that the highest inflammatory response occurred 6 hours after PHA injection (results not shown) and thus we based our comparisons only for values obtained in this period. Because we found higher inflammatory response only after the second injection of PHA (see Results), we estimated the fractional change in *SI* from day 1 to day 10 in antibiotic-treated and control bats as

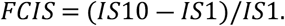

### Assessment of the adaptive humoral response

We measured IgG concentration in plasma collected from each bat 24 hours before the 1^st^ injection and 24 hours after the 3^rd^ injection of PHA. Plasma was separated from whole blood after being centrifuged; the samples were immediately placed in liquid nitrogen and then transferred to an ultra-low temperature freezer until they were analyzed in the laboratory. We measured IgG from 12 and 14 individuals in the control and the experimental groups, respectively, because were not able to collect enough plasma from some individuals. To determine the IgG concentration, we performed a 1:30,000 dilution of blood plasma, according to protocols established for the species [30, 31] in a 0.1M NaHCO3 solution, pH 9.6 in a 96-well microplate (Corning 3596, Kennebunk, ME, USA). Plasma was then incubated for 18 hours at 4°C. After incubation, the plate was washed three times with 200 µl of PBS-Tween-20 (0.05%) to remove excess diluted plasma, then we added 50 µL of bovine serum albumin (BSA, RMBIO Bovine Serum Albumin IgG free, USA) diluted 1% in PBS (0.01 M) to block non-specific binding and it was incubated for 2 hours at room temperature. We then washed the plate three times with PBS-Tween-20 (0.05%) and then added 50 µL of Goat polyclonal anti-bat IgG conjugate with HRP (Novus Biologicals, NB7238; 1:10000 in PBS-BSA 1% solution). We continued incubating for two hours and then washed 3 times in PBS-Tween-20 (0.05%). Finally, we added 50 µL of a developer solution (SIGMAFAST OPD p9187, Sigma Aldrich, Mexico) and after 5 minutes we stopped the reaction with 100 µL of H_2_SO_4_ (0.01%). We measured the absorbance of each well at 490 nm per sample in triplicate. Antibody concentration is directly proportional to the absorption [32]; therefore, we estimated IgG concentration as the mean optical density (OD) of the 3 replicates for each sample. We estimated the fractional change in IgG concentration from day 1 to day 15 in antibiotic-treated and control bats as

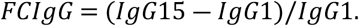

### Analyses of data

Data were checked for normality and analyzed with parametric or non-parametric tests accordingly. We compared *SI* values after the first and second PHA injection using repeated measures analyses of variance (RM-ANOVA) for antibiotic-treated and for control bats. We compared the *FCSI*of antibiotic-treated and control bats using a Mann-Whitney U test. We compared IgG levels after the first and third PHA injections using RM-ANOVA for antibiotic-treated and for control bats. We compared the *FCIgG*of antibiotic-treated and control bats using a Mann-Whitney U test. The analyzes were conducted using the statistical program PAST 4.06b [33], and we used a fiducial level of 0.05 for significance of the results.

## Results

### Inflammatory response

The *SI* was higher after the second injection with respect to the first injection in control bats (*F*_1, 22_ = 4.992, p = 0.040; Fig. 1A), but it did not differ among injection periods for bats treated with antibiotic (F_1, 28_ = 0.561, p = 0.466; Fig. 1B). The *FCIS* were significantly different between treatments (*U* = 54.0, p = 0.02): the *SI* increased to a higher rate after the second PHA injection in control bats (n =14) than in bats treated with antibiotic (n =15; Fig. 1C).

**Figure 1.**
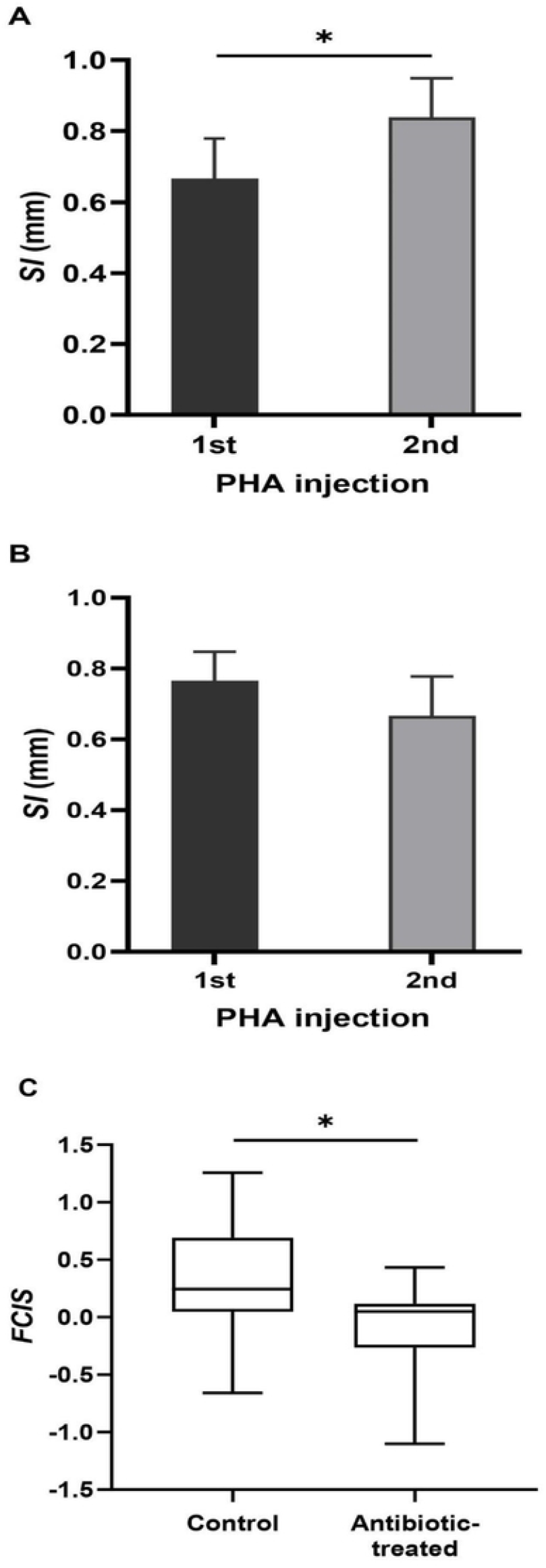
Inflammatory response after the repeated application of phytohemagglutinin (PHA) in the footpad of common vampire bats (*Desmodus rotundus*). We compared the swelling index (*SI*; mean ± SE) following a first and a second injections of PHA in control bats (A) and bats treated with an antibiotic (B), and the fractional change in *SI (FCSI*; median, 25% interquartile range, minimum-maximum range) after the second PHA injection between control and antibiotic-treated bats (C). *p ≤ 0.05.

### IgG levels

IgG concentration values were significantly higher following the third PHA injection than before the first injection of PHA in control bats (*F*_1, 22_ = 4.975, p = 0.047; Fig.2A), but they were not significantly different between injection periods in antibiotic-treated bats (*F*_1, 26_ = 3.288, p = 0.092; Fig 2B). The *FCIgG* was significantly different between treatments (*U* = 28.00, p =0.004): IgG concentration of control bats (n = 12) increased at a higher rate than that of bats treated with antibiotic (n = 14; Fig. 2C).

**Figure 2.**
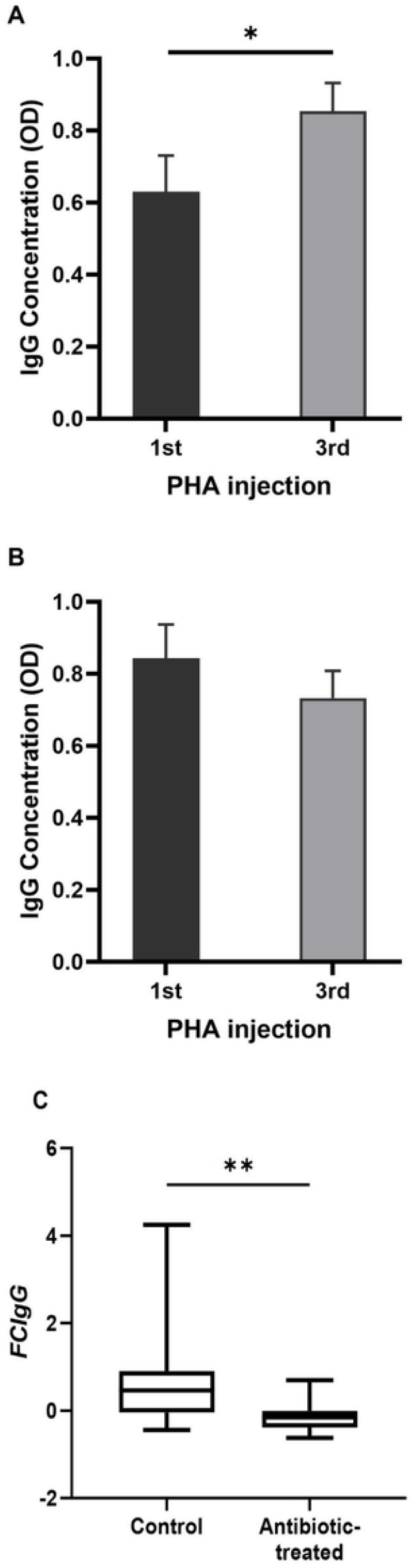
Immunoglobulin G (IgG) concentration in serum of common vampire bats (*Desmodus rotundus*) before and after phytohemagglutinin (PHA) application. We compared IgG (mean ± SE) before a first and after a third PHA injections in control bats (A) and bats treated with antibiotic (B), and the fractional change in IgG concentration (*FCIG*; median, 25% interquartile range, minimum-maximum range) after the third PHA injection between control and antibiotic-treated bats (C). IgG concentration is the mean optical density (OD) of the 3 replicates for each sample. *p ≤ 0.05, **p ≤ 0.01.

## Discussion

### Cellular response

According to our predictions, antibiotic-exposed vampire bats exhibited weaker inflammatory response to the repeated PHA treatment. A second injection of PHA in vampire bats that were not treated with antibiotic induced swelling that was ∼30% larger than that after the first injection, whereas swelling after the second injection in antibiotic-treated bats was ∼10% lower than after the first injection. A secondary injection of PHA in antibiotic-free vampire bats led to a similar response than that observed in blue-footed booby (*Sulane bouxii*; [22]), several species of parrots [21], and toads (*Rhinella marina*; [23]. The functional significance of PHA-induced swelling in bats has been examined previously but not after repeated PHA injections. A single PHA injection increases the presence of leukocytes in Brazilian free-tailed bats (*Tadarida brasiliensis*), but the magnitude of skin swelling has a marginally positive correlation with basophil abundance and no correlation with the abundance of other leukocytes (lymphocytes, neutrophils, macrophages, and eosinophils; [34]).

Although there are no previous studies specifically testing the effect of antibiotics on local swelling after PHA exposure, our findings are in line with other studies that explored their effect on cellular immune components. For instance, application of chloramphenicol and florfenicol decreases the proliferation of B and T cells in common carps [11], T cells (Th1 and Th17) decrease in pregnant house mice after being treated with a mixture of neomycin, bacitracin and pimaricin [35], an application of several antibiotics suppresses DNA synthesis [36] and mitogenesis [37] of human leukocytes stimulated with PHA. In addition to its bactericidal properties, clindamycin modulates the production of inflammatory cytokines [38]. The lower inflammatory response that we found in clindamycin-treated vampires might also be a response to a lower secretion of inflammatory cytokines by PHA-stimulated lymphocytes, similarly to the effects of other pharmacological products on PHA-derived inflammation [39]. Further work should test if the reduced inflammatory response observed in clindamycin-treated vampires after repeated PHA injections is due to reduced production of leukocytes, modulation of inflammatory cytokines, or both.

### Humoral response

PHA stimulates the growth of blood lymphocytes triggering the synthesis of immunoglobulins [40]. For instance, PHA stimulates the production of IgG and IgM in peripheral blood lymphocytes of domestic dogs [41], and it increases serum levels of IgE [42] and IgG [43] in house mice. Accordingly, we found an increase of IgG serum levels following three consecutive PHA injections but it occurred only in vampires that did not receive the antibiotic. The median value of IgG serum levels of control individuals after three PHA injections increased ∼90% over pre-injection values, whereas this value was ∼15% lower than pre-injection values in vampires treated with antibiotic. Although not specifically tested after an immune challenge as in our study, the detrimental effect of antibiotics on immunoglobulin production has been previously reported mostly in humans and domesticated animals. The immunoglobulin production decreases after the application of florfenicol [44] and ciprofloxacin [45] in domestic chicken (*Gallus domesticus*), of oxytetracycline in common carps [9], of amoxicillin and clavulanate in humans [10, 46], and of a mixture of amoxicillin, doxycycline and tilmicosin in domestic pigs [10].

### Concluding remarks and future directions

Due to their highly specialized feeding habits, the common vampire bat is reservoir of different bacteria with zoonotic potential [47], and they are potentially exposed to the consumption of antibiotics when feeding on livestock. We found that antibiotic exposure negatively affected the performance of the acquired immune system of vampire bats. Previous non-experimental evidence with vampire bats also points to the potential pernicious effect of cattle-feeding on their immune performance. Common vampire bats captured in sites close to livestock show lower expression of acquired immune response (IgG levels, and lymphocyte proportion) than those that feed mostly on wildlife [31]. Exposure of wildlife to antibiotics might affect their gut microbiota composition which in turn might impact the performance of their immune system [48, 49]. For instance, dysbiosis might reduce the presence of some members of the intestinal microbiota that induce the synthesis of systemic IgG [47, 50]. Further experimental work with vampire bats should determine the effects of antibiotics on their gut microbiota and test if their modification is linked to changes in their immune response. Observational work with common vampire bats shows that changes in core bacteria composition were associated to their use of livestock as food [47]. Our study adds to previous evidence pointing to the negative effect that exposure to anthropogenic chemicals (e.g., mercury) generates in the capacity of bats to maintain a healthy immune system [30, 51].

## Acknowledgments

This article is part of the requirements for I.C.-C. to obtain a PhD degree at the Universidad Autónoma de Tlaxcala. We thank Marco Tulio Solano De la Cruz (Unidad de Genética Molecular of the Instituto de Ecología-Universidad Nacional Autónoma de México) for technical support during the immunoglobulin laboratory analyses.

## Author Contributions

**Conceptualization:** Iván Cabrera-Campos, L. Gerardo Herrera M.

**Data curation:** Iván Cabrera-Campos, L. Gerardo Herrera M.

**Data analyses:** Iván Cabrera-Campos, L. Gerardo Herrera M.

**Funding acquisition:** L. Gerardo Herrera M.

**Investigation:** Iván Cabrera-Campos, Rafael Ávila Flores, David Alfonso Rivera-Ruiz, L. Gerardo Herrera M.

**Project administration:** L. Gerardo Herrera M.

**Resources:** L. Gerardo Herrera M., Rafael Ávila Flores.

**Supervision:** L. Gerardo Herrera M., Rafael Ávila Flores.

**Writing – original draft**: Iván Cabrera-Campos, L. Gerardo Herrera M.

**Writing – review & editing:** Rafael Ávila Flores, David Alfonso Rivera-Ruiz

## References

1. aus der Beek T, Weber FA, Bergmann A, Hickmann S, Ebert I, Hein A, et al. Pharmaceuticals in the environment—Global occurrences and perspectives. Environmental Toxicology and Chemistry. 2016;35:823–35.

2. Khan AH, Aziz HA, Khan NA, Hasan MA, Ahmed S, Farooqi IH, et al. Impact, disease outbreak and the eco-hazards associated with pharmaceutical residues: a critical review. International Journal of Environmental Science and Technology. 2021:1–12.

3. Oaks JL, Gilbert M, Virani MZ, Watson RT, Meteyer CU, Rideout BA, et al. Diclofenac residues as the cause of vulture population decline in Pakistan. Nature. 2004;427:630–3.

4. Laborda P, Sanz-García F, Ochoa-Sánchez LE, Gil-Gil T, Hernando-Amado S, Martínez JL. Wildlife and antibiotic resistance. Frontiers in Cellular and Infection Microbiology. 2022;12:568.

5. Saaristo M, Brodin T, Balshine S, Bertram MG, Brooks BW, Ehlman SM, et al. Direct and indirect effects of chemical contaminants on the behaviour, ecology and evolution of wildlife. Proceedings of the Royal Society B. 2018;285:20181297.

6. Polianciuc S, Gurzău A, Kiss B, Ştefan M-G, Loghin F. Antibiotics in the environment: causes and consequences. Medicine and Pharmacy Reports. 2020;93.

7. Ogawa M, Goto S, Ishikawa F, Kimura I. Effects of various antibiotics on antibody-producing cells of mouse spleen. Chemotherapy. 1986;32:464–7.

8. Furuhama K, Benson W, Knowles J, Roberts DW. Immunotoxicity of cephalosporins in mice. Chemotherapy. 1993;39:278–85.

9. Rijkers GT, Van Oosterom R, Van Muiswinkel WB. The immune system of cyprinid fish. Oxytetracycline and the regulation of humoral immunity in carp (Cyprinus carpio). Veterinary Immunology and Immunopathology. 1981;2:281–90.

10. Bosi P, Merialdi G, Scandurra S, Messori S, Bardasi L, Nisi I, et al. Feed supplemented with 3 different antibiotics improved food intake and decreased the activation of the humoral immune response in healthy weaned pigs but had differing effects on intestinal microbiota. Journal of Animal Science. 2011;89:4043–53.

11. Sieroslawska. A., M. Studnicka. A. K. Siwicki. A. Bownik. A. Rymuszka. J. Sionka. Antibiotics and cell-mediated immunity in fish -in vitro study. Acta Veterinaria Brno 1998.67: 329–334.

12. Marshall BM, Levy SB. Food animals and antimicrobials: impacts on human health. Clinical Microbiology Reviews. 2011;24:718–33.

13. Brown N, Escobar LE. A review of the diet of the common vampire bat (Desmodus rotundus) in the context of anthropogenic change. Mammalian Biology. 2023:1–21.

14. Mendoza-Sáenz VH, Saldaña-Vázquez RA, Navarrete-Gutiérrez D, Kraker-Castañeda C, Ávila-Flores R, Jiménez-Ferrer G. Reducing conflict between the common vampire bat Desmodus rotundus and cattle ranching in Neotropical landscapes. Mammal Review. 2023;53:72–83.

15. Lafferty KD, Kuris AM. Trophic strategies, animal diversity and body size. Trends in Ecology and Evolution. 2002;17:507–13.

16. Benavides JA, Godreuil S, Opazo-Capurro A, Mahamat OO, Falcon N, Oravcova K, et al. Long-term maintenance of multidrug-resistant Escherichia coli carried by vampire bats and shared with livestock in Peru. Science of the Total Environment.2022;810:152045.

17. Carrillo Gaeta N, Cavalcante Brito JE, Nunes Batista JM, Gagete Veríssimo de Mello B, Dias RA, Heinemann MB. Bats Are carriers of antimicrobial-resistant Staphylococcaceae in their skin. Antibiotics. 2023;12:331.

18. Martin LB, Han P, Lewittes J, Kuhlman JR, Klasing KC, Wikelski M. Phytohemagglutinin-induced skin swelling in birds: histological support for a classic immunoecological technique. Functional Ecology. 2006;20:290–9.

19. Vinkler M, Bainová H, Albrecht T. Functional analysis of the skin-swelling response to phytohaemagglutinin. Functional Ecology. 2010;24:1081–6.

20. Vinkler M, Svobodová J, Gabrielová B, Bainová H, Bryjová A. Cytokine expression in phytohaemagglutinin-induced skin inflammation in a galliform bird. Journal of Avian Biology. 2014;45:43–50.

21. Tella JL, Lemus JA, Carrete M, Blanco G. The PHA test reflects acquired T-cell mediated immunocompetence in birds. PLOS ONE. 2008;3:e3295.

22. Santiago-Quesada F, Albano N, Castillo-Guerrero JA, Fernández G, González-Medina E, Sánchez-Guzmán JM. Secondary phytohaemagglutinin (PHA) swelling response is a good indicator of T-cell-mediated immunity in free-living birds. Ibis. 2015;157:767–73.

23. Brown GP, Shilton CM, Shine R. Measuring amphibian immunocompetence: validation of the phytohemagglutinin skin-swelling assay in the cane toad, Rhinella marina. Methods in Ecology and Evolution. 2011;2:341–8.

24. Ávila-Flores R, Bolaina-Badal AL, Gallegos-Ruiz A, Sánchez-Gómez WS. Use of linear features by the common vampire bat (Desmodus rotundus) in a tropical cattle-ranching landscape. Therya. 2019;10:229–34.

25. Lollar A. Standards and medical management for captive insectivorous bats. Bat World Sanctuary. 2010;1:1–204.

26. Sikes RS, the Animal Care and Use Committee of the American Society of Mammalogists. 2016 Guidelines of the American Society of Mammalogists for the use of wild mammals in research and education. Journal of Mammalogy. 2016;97:663–88.

27. Otalora-Ardila A, Herrera M LG, Flores-Martínez JJ, Welch Jr KC. Metabolic cost of the activation of immune response in the fish-eating myotis (Myotis vivesi): the effects of inflammation and the acute phase response. PLOS ONE. 2016;11:e0164938.

28. Allen LC, Turmelle AS, Mendonça MT, Navara KJ, Kunz TH, McCracken GF. Roosting ecology and variation in adaptive and innate immune system function in the Brazilian free-tailed bat (Tadarida brasiliensis). Journal of Comparative Physiology B. 2009;179:315–23.

29. Smits JE, Bortolotti GR, Tella JL. Simplifying the phytohaemagglutinin skin-testing technique in studies of avian immunocompetence. Functional Ecology. 1999;13:567–72.

30. Becker DJ, Chumchal MM, Bentz AB, Platt SG, Czirják GÁ, Rainwater TR, et al. Predictors and immunological correlates of sublethal mercury exposure in vampire bats. Royal Society Open Science. 2017;4:170073.

31. Becker DJ, Czirják GÁ, Volokhov DV, Bentz AB, Carrera JE, Camus MS, et al. Livestock abundance predicts vampire bat demography, immune profiles and bacterial infection risk. Philosophical Transactions of the Royal Society B: Biological Sciences. 2018;373:20170089.

32. Schneeberger K, Courtiol A, Czirják GÁ, Voigt CC. Immune profile predicts survival and reflects senescence in a small, long-lived mammal, the greater sac-winged bat (Saccopteryx bilineata). PLOS ONE. 2014;9:e108268.

33. Hammer O, Harper D, Ryan P. PAST: Paleontological statistics software package for education and data analysis. Palaeontologia Electronica. 2001;4:1–9.

34. Turmelle AS, Ellison JA, Mendonça MT, McCracken GF. Histological assessment of cellular immune response to the phytohemagglutinin skin test in Brazilian free-tailed bats (Tadarida brasiliensis). Journal of Comparative Physiology B. 2010;180:1155–64.

35. Faas MM, Liu Y, Wekema L, Weiss GA, van Loo-Bouwman CA, Silva Lagos L. The effect of antibiotics treatment on the maternal immune response and gut microbiome in pregnant and non-pregnant mice. Nutrients. 2023;15:2723.

36. Munster AM, Loadholdt CB, Leary AG, Barnes MA. The effect of antibiotics on cell-mediated immunity. Surgery. 1977;81:692–5.

37. Rachkov SM, Bekbergenov BM, Korolev PN, Glezer GA. Effect of antibiotics on human lymphocyte mitogenesis. Antibiotiki. 1981;26:55–8.

38. Nakano T, Hiramatsu K, Kishi K, Hirata N, Kadota J-i, Nasu M. Clindamycin modulates inflammatory-cytokine induction in lipopolysaccharide-stimulated mouse peritoneal macrophages. Antimicrobial Agents and Chemotherapy. 2003;47:363–7.

39. Askari VR, Rahimi VB, Zargarani R, Ghodsi R, Boskabady M, Boskabady MH. Anti-oxidant and anti-inflammatory effects of auraptene on phytohemagglutinin (PHA)-induced inflammation in human lymphocytes. Pharmacological Reports. 2021;73:154–62.

40. Greaves MF, Roitt IM. The effect of phytohaemagglutinin and other lymphocyte mitogens on immunoglobulin synthesis by human peripheral blood lymphocytes in vitro. Clinical and Experimental Immunology. 1968;3:393.

41. Letwin BW, Quimby FW. Effects of concanavalin A, phytohemagglutinin, pokeweed mitogen, and lipopolysaccharide on the replication and immunoglobulin synthesis by canine peripheral blood lymphocytes in vitro. Immunology Letters. 1987;14:79–85.

42. Astorquiza MI, Cisternas C, Leal X. Sex-dependent differences in the IgE response modulated by phytohemagglutinin. Immunology Letters. 1987;16:27–30.

43. Vogel SN, Roberson BS. Phytohemagglutinin stimulation of enhanced immunoglobulin G production in mice inoculated with type III pneumococcal polysaccharide. Infection and Immunity. 1978;22:901–7.

44. Cao J, Li K, Lu X, Zhao Y. Effects of florfenicol and chromium (III) on humoral immune response in chicks. Asian-australasian Journal of Animal Sciences. 2004;17:366–70.

45. Niyogi D, Bhowmik MK, Chatterjee C. Effect of ciprofloxacin on humoral immune response in broiler chicken. Indian Journal of Animal Health. 2000;39:63–4.

46. Dufour V, Millon L, Faucher J-F, Bard E, Robinet E, Piarroux R, et al. Effects of a short-course of amoxicillin/clavulanic acid on systemic and mucosal immunity in healthy adult humans. International Immunopharmacology. 2005;5:917–28.

47. Ingala MR, Becker DJ, Bak Holm J, Kristiansen K, Simmons NB. Habitat fragmentation is associated with dietary shifts and microbiota variability in common vampire bats. Ecology and Evolution. 2019;9:6508–23.

48. Thomason CA, Mullen N, Belden LK, May M, Hawley DM. Resident microbiome disruption with antibiotics enhances virulence of a colonizing pathogen. Scientific Reports. 2017;7:16177.

49. Bravo M, Combes T, Martinez FO, Risco D, Gonçalves P, Garcia-Jimenez WL, et al. Wildlife Symbiotic Bacteria Are Indicators of the Health Status of the Host and Its Ecosystem. Applied and Environmental Microbiology. 2022;88:e01385–21.

50. Zeng MY, Cisalpino D, Varadarajan S, Hellman J, Warren HS, Cascalho M, et al. Gut microbiota-induced immunoglobulin G controls systemic infection by symbiotic bacteria and pathogens. Immunity. 2016;44:647–58.

51. Costantini D, Czirják GÁ, Bustamante P, Bumrungsri S, Voigt CC. Impacts of land use on an insectivorous tropical bat: The importance of mercury, physio-immunology and trophic position. Science of the Total Environment. 2019;671:1077–85.

